# Unsupervised Learning of Temporal Features for Word Categorization in a Spiking Neural Network Model of the Auditory Brain

**DOI:** 10.1101/059840

**Authors:** Irina Higgins, Simon Stringer, Jan Schnupp

**Affiliations:** Department of Experimental Psychology, University of Oxford, Oxford, England; Department of Physiology, Anatomy and Genetics (DPAG), University of Oxford, Oxford, England

## Abstract

The nature of the code used in the auditory cortex to represent complex auditory stimuli, such as naturally spoken words, remains a matter of debate. Here we argue that such representations are encoded by stable spatio-temporal patterns of firing within cell assemblies known as polychronous groups, or PGs. We develop a physiologically grounded, unsupervised spiking neural network model of the auditory brain with local, biologically realistic, spike-time dependent plasticity (STDP) learning, and show that the plastic cortical layers of the network develop PGs which convey substantially more information about the speaker independent identity of two naturally spoken word stimuli than does rate encoding that ignores the precise spike timings. We furthermore demonstrate that such informative PGs can only develop if the input spatio-temporal spike patterns to the plastic cortical areas of the model are relatively stable.

**Author Summary:** Currently we still do not know how the auditory cortex encodes the identity of complex auditory objects, such as words, given the great variability in the raw auditory waves that correspond to the different pronunciations of the same word by different speakers. Here we argue for temporal information encoding within neural cell assemblies for representing auditory objects. Unlike the more traditionally accepted rate encoding, temporal encoding takes into account the precise relative timing of spikes across a population of neurons. We provide support for our hypothesis by building a neurophysiologically grounded spiking neural network model of the auditory brain with a biologically plausible learning mechanism. We show that the model learns to differentiate between naturally spoken digits “one” and “two” pronounced by numerous speakers in a speaker-independent manner through simple unsupervised exposure to the words. Our simulations demonstrate that temporal encoding contains significantly more information about the two words than rate encoding. We also show that such learning depends on the presence of stable patterns of firing in the input to the cortical areas of the model that are performing the learning.

## Introduction

The nature of the neural code used by the auditory brain to represent complex auditory stimuli, such as naturally spoken words, remains uncertain [1,2]. A variety of spike rate and spike timing coding schemes are being debated. Rate encoding presumes that the identity of an auditory stimulus is encoded by the average firing rate of a subset of neurons, but the precise timing of individual spikes is irrelevant. Temporal encoding suggests that different auditory stimuli are represented by spatio-temporal patterns of spiking activity within populations of neurons, where the relative timing of the spikes is part of the representation.

A widely held view of the auditory pathway is that temporal encoding plays a major role in the early subcortical areas, but becomes increasingly less important in the midbrain and the cortical areas [3]. Here we build on existing theories of learning in spiking neural networks [4–7] to argue that temporal coding may have a crucial role to play in the auditory cortex. In particular, we argue that the basic information encoding units for representing complex auditory stimuli, such as naturally spoken words, in the auditory cortex are spatio-temporal patterns of firing within cell assemblies called polychronous groups (PGs) [6].

Our hypothesis is evaluated using a biologically inspired hierarchical spiking neural network model of the auditory brain comprising of the auditory nerve (AN), cochlear nucleus (CN), inferior colliculus (IC), and auditory cortex (CX) stages, where the last CX stage includes primary (A1) and “higher order” (Belt) cortex (Fig. 1). Using only biologically plausible local spike-time dependent plasticity (STDP) learning [8], our auditory brain model is trained to discriminate between two naturally spoken words, “one” and “two”, through unsupervised exposure to the stimuli. In order to succeed on the task, the model has to learn how to cope with the great variability of the acoustic waveforms of these sounds when pronounced by many different speakers (TIDIGITS database [9]).

We show that stable spatio-temporal patterns of firing (PGs) spontaneously emerge within the CX stage of the model, and that they are significantly more informative of the auditory object identity than an alternative rate coded information encoding scheme that disregards the precise spike timing information. Furthermore, our results show that such PG-based learning in the plastic cortical stages of the model relies on relatively stable spatio-temporal input patterns. Due to the stochasticity of spiking times in the AN, if the AN spike patterns are fed directly to the plastic cortical areas, these spike patterns are not stable enough for the emergence of informative PGs in the cortical areas [10]. Hence, the particular subcortical circuitry of the CN and IC is necessary to reduce the high levels of noise known to exist in the AN and re-introduce stability within the firing patterns that serve as input to the plastic cortical areas of the model, thus enabling PG-based learning to emerge in the plastic CX [10].

**Fig 1.**
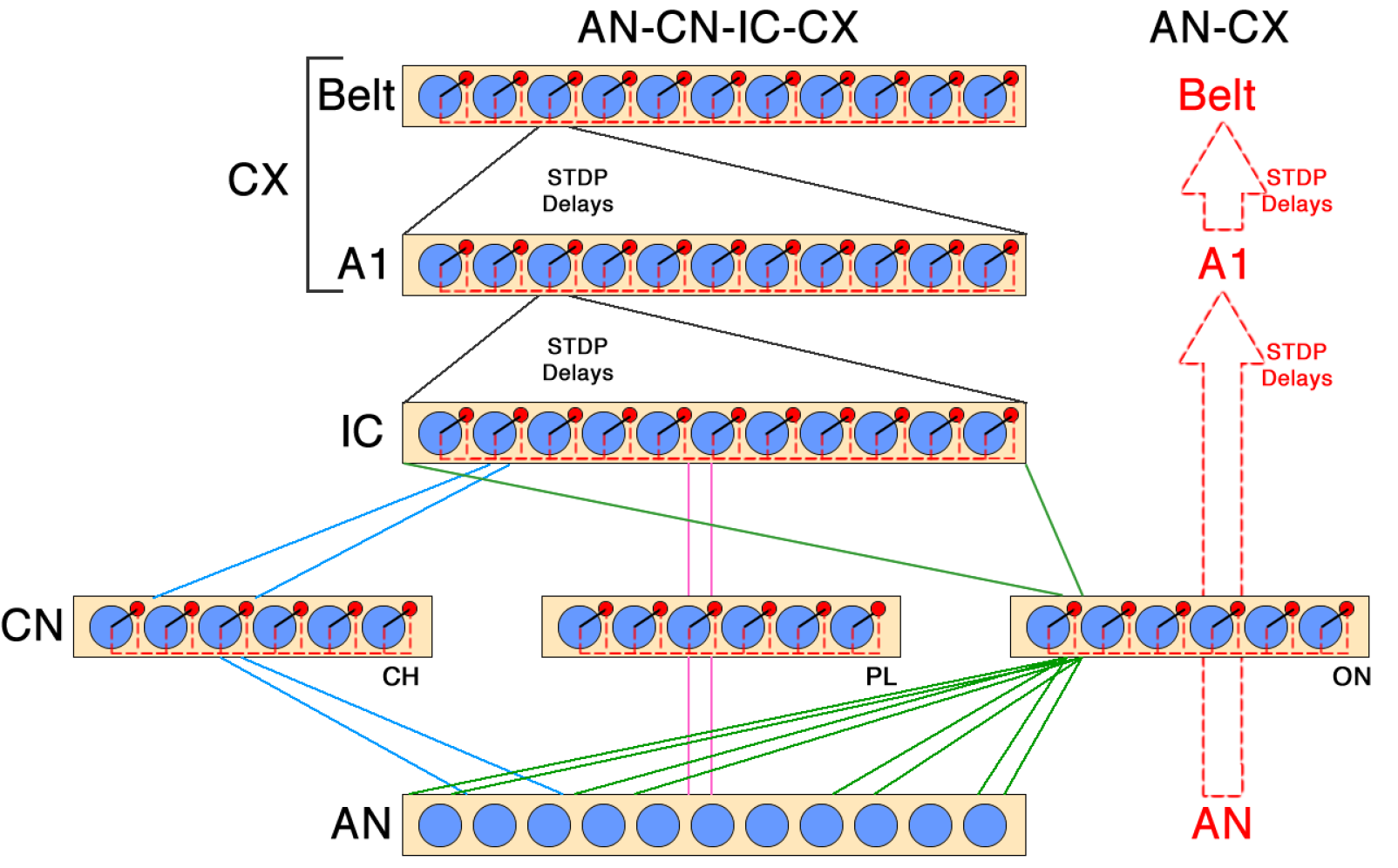
Schematic representation of the full AN-CN-IC-CX and reduced AN-CX models. In the AN-CX model, direct plastic connections from the AN to the A1 replace CN and IC layers. Blue circles are excitatory, red circles inhibitory cells.

The main contributions of our paper are two-fold: 1) we provide simulation evidence to argue for spatio-temporal information encoding using PGs in the auditory cortex; and 2) we demonstrate that PG-based learning in the plastic auditory cortex relies on the relative stability of the input spatio-temporal firing patterns.

## Materials and Methods

### Learning mechanisms

In order to form speaker independent representations of different words, the auditory brain has to be able to respond in a manner that discriminates between different words but not between different exemplars of the same word. This is a challenging task, given the great variability in the raw auditory waves corresponding to the same word due to differences in pronunciation both within and between speakers. This input variability is further compounded by the stochasticity present in the firing patterns generated at the first neural stage of auditory processing, the AN. How can the brain discover the statistical regularities differentiating various words in such noisy inputs? We believe an answer to this question can be found in a number of independent yet overlapping theories describing how spiking neural networks with local STDP learning may discover and amplify the statistical regularities in temporal input patterns [4-7].

**Learning to extract repeating patterns from noise**: The first relevant idea was described by [5], who showed that a single spiking neuron can learn to pick out a repeating spatio-temporal pattern of firing from statistically identical noise using STDP learning (see Fig. 2A). This effect depends on the particular biologically realistic STDP learning configuration with stronger long term depression (LTD) (parameters *α_d_* and *τ*_d_ in our models, see Tbl. 1) compared to long term potentiation (LTP) (parameters *α_p_* and *τ*_p_ in our models, see Tbl. 1). Such STDP configuration results in the overall weakening of the feedforward connections (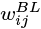 in our models, see Tbl. 1) to the output neuron due to the random input firing in response to noise. This is true for all connections apart from those originating from the input cells involved in the repeating pattern, which instead get strengthened due to repeating LTP every time the pattern is presented. Such simple circuits are able to cope with variable or degraded inputs while maintaining stable, informative output representations. For example, they may learn to cope with the presence of temporal jitter in the input on the order of a few milliseconds, additive Poisson noise of up to 10 Hz, or loss of up to half of the neurons participating in the input repeating pattern [5].

**Table 1.**
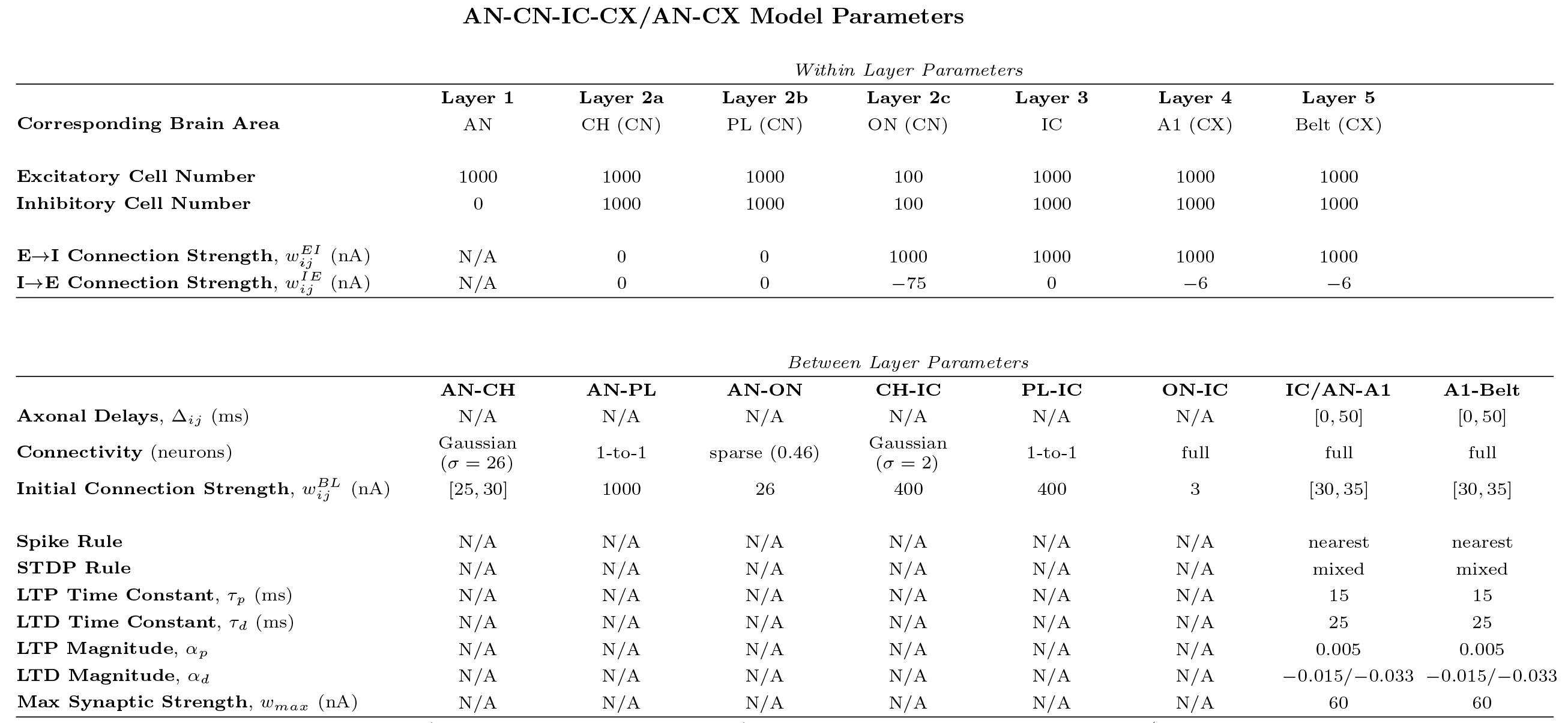
Parameters used for the full AN-CN-IC-CX and reduced AN-CX models of the auditory brain (where these differ for the LTD magnitude (*α_d_*), the AN-CX parameters are given as second values following a slash). AN – auditory nerve; CN – cochlear nucleus with three subpopulations of cells discriminated based on their connectivity: chopper (CH), primary-like (PL) and onset (ON); IC – inferior colliculus; A1 – primary auditory cortex; Belt – belt area of the auditory cortex. The parameters were found to be optimal using a grid search heuristic on a two vowel recognition task (see [10] for details). The sparse connectivity parameter for the AN to ON connections defines the proportion of dead synapses between these two layers.

**Extending the memory capacity and temporal receptive fields for repeating pattern learning**: The output neuron described by [5] can learn one short spatio-temporal pattern of firing (see Fig. 2A). The learning depends on firing co-occurences within the pattern that lie within the neuron’s temporal integration window (on the order of a few milliseconds). Such a setup in its original form therefore has limited value for learning longer spatio-temporal patterns, such as words, that can last on the order of hundreds of milliseconds. This shortcoming can be partially averted by the addition of randomly initialised conduction delays (Δ_*ij*_ in our models, see Tbl. 1) for the feed-forward connections (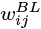 in our models, see Tbl. 1). The conduction delays would extend the temporal range of coincidences that each output neuron can detect (see Fig. 2B).

**Fig 2.**
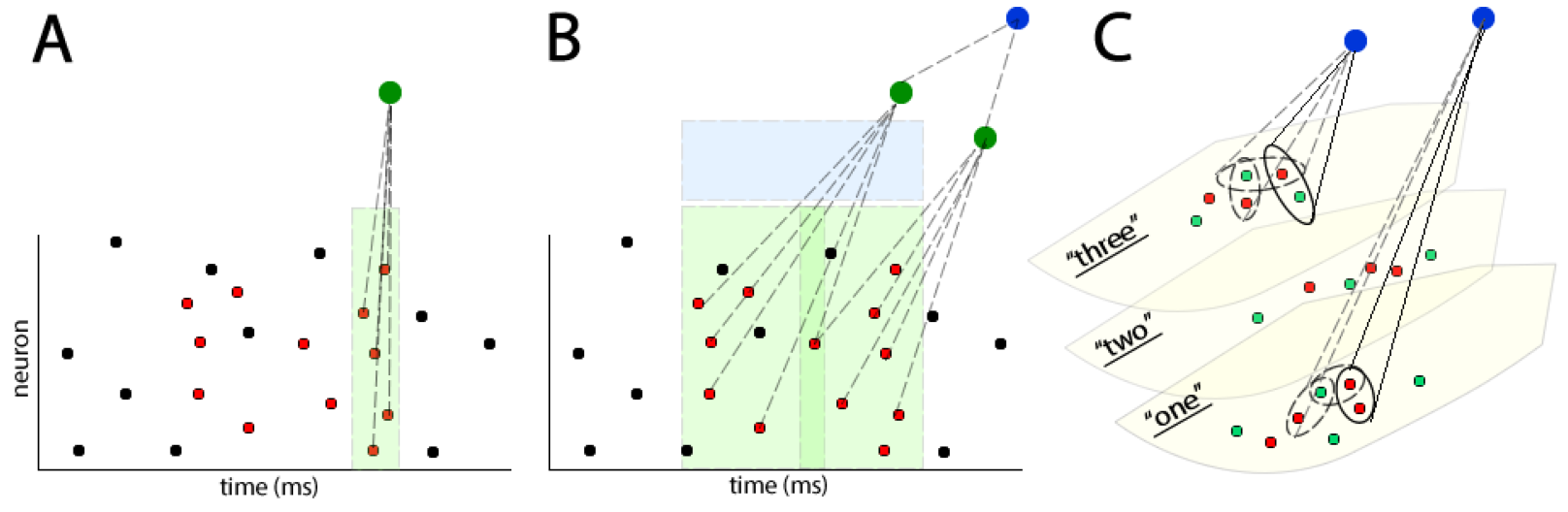
**A**: single output neuron (green) can learn to pick out a repeating spatio-temporal pattern of firing (red) out of statistically identical noise (black) [5]. In this example the output neuron relies on concurrent input from at least five input neurons in order to fire. Due to instantaneous axonal conductances, the neuron has a very narrow temporal integration window of a few milliseconds (shown in light green). **B**: If random axonal conduction delays (Δ_*ij*_ in our models, see Tbl. 1) are added for the feedforward connections (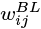 in our models, see Tbl. 1), each neuron in the next stage of the model (green) becomes sensitive to a particular pattern of firing in the input. Axonal conduction delays extend temporal integration windows of output neurons (shown in light green). Adding extra output layers with random distributions of axonal delays creates a hierarchy of pattern learning neurons. Neurons at the end of such a hierarchy (blue) have the largest temporal integration windows (shown in light blue) **C**: different pronunciations of words “one”, “two”, and “three” by two different speakers (red and blue dots respectively) lie on different low dimensional manifolds. Polychronous groups (PGs) [6] in the auditory cortex (blue circles) can learn to become sensitive to similar pronunciations of one preferred word (solid ovals). Continuous transformation learning [4] extends the sensitivity of PGs to more different pronunciations of the preferred word (dashed ovals), while maintaining the selectivity of PGs to exemplars of one word only.

If an output neuron receives input spikes through connections with randomly initialised delays, it will only fire if the right input neurons fire in the right temporal order that matches the delay lines. Only then would their spikes arrive at the output neuron coincidentally and depolarise it enough to fire. Since different output neurons will have different axonal delays initialised for their afferent connections, they will be sensitive to different input patterns of firing. Hence, the addition of extra output neurons with different randomly initialised delays would introduce heterogeneity in the types of spatio-temporal patterns the output layer as a whole can learn (see Fig. 2B). Such heterogeneity would allow the feedforward network to organise its firing into a hierarchy of “polychronous groups” (PGs) [6]. PGs are stable spatio-temporal patterns of firing, where neurons within a layer “exhibit reproducible time-locked but not synchronous firing patterns with millisecond precision” [6]. Each neuron can be part of numerous PGs, thus increasing network memory capacity [6]. The idea is that PGs in each layer will be sensitive to particular parts of repeating spatio-temporal patterns that are characteristic of a particular stimulus class. Throughout the hierarchy of the network, PGs will emerge that are more invariant to the different variations of their preferred pattern and have longer temporal receptive fields. The details of such PG-based learning is discussed next.

The nature of learnt PGs is shaped by the interplay between delay lines, STDP learning and stimulus structure. A random distribution of conduction delays sets up a repertoire of PGs in a network as described above. When a stimulus is presented to the network, the resulting input spatio-temporal firing patterns are propagated through a set of connections with random delays. If that set of connections is large then it may contain subsets of connections with delays which happen to match the characteristic spatio-temporal firing patterns in the input in a manner that allows the input spikes to converge synchronously on a receiving neuron. This receiving neuron thereby receives super-threshold activation, and its connections to the input spatio-temporal pattern are strengthened by STDP learning.

In this manner, different output layer neurons become sensitized to the characteristic activity of different patterns of firing in the input layer. The temporal structure of the input stimuli may cause the output layer neurons themselves to generate reproducible spatio-temporal firing patterns, giving rise to “higher order” PGs, which may in turn be learned by the next layer in the network. Such a feedforward hierarchy could take advantage of cumulative delays over several layers of connectivity, enabling PGs to discover regularities in the temporal structure of input stimuli over an ever wider temporal scale (see Fig. 2B).

**Extending robustness to pattern variability**: While building on the setup described by [5], PG-based learning is still not quite sufficient to enable a feedforward spiking neural network to form speaker independent representations of naturally spoken words, because it is unable to cope with the high degree of pronunciation variability. To tackle this we introduce the last relevant concept: the Continuous Transformation (CT) learning principle [4]. CT learning is a mechanism originally developed to describe geometric transform invariant visual object recognition in a rate-coded neural network model of the ventral visual stream [11]. It takes advantage of the fact that when visual objects undergo smooth transformations, such as rotations, translations or scalings, the nearest neighbours of the resulting projections into the two-dimensional retinal input space have a high degree of overlap or correlation. CT learning binds these similar input patterns together into an invariant representation of that object (or an object orbit according to [12]) and maps them onto the same subset of higher stage neurons.

While originally conceived around learning transform invariant representations of visual objects, CT learning generalizes to other modalities where the nearest neighbours of different exemplars of a given stimulus class strongly overlap in the high dimensional raw sensory input space. For example, different pronunciations of the same word might form a low-dimensional manifold for that word, and different words might form different manifolds (see Fig. 2C). Two different pronunciations of one word may have highly overlapping AN spatio-temporal firing rasters, and hence be nearest neighbours on the corresponding low-dimensional manifold for that word. By chance, a PG may be sensitive to a particular region of that word manifold. CT learning would then “expand” the span of such a PG to a more extensive region of the manifold, while preserving the selectivity of the PG to its preferred word manifold only. In this way a hierarchy of PGs would emerge through learning in a feed-forward spiking neural network, whereby PGs higher up in the hierarchy would learn to respond to an increasing number of pronunciations of the same word, while ignoring the pronunciations of all other words.

### Stimuli

Recordings of two pronunciations of the digits “one” and “two” from each of 94 native American English speakers served as stimuli (TIDIGITS database [9]). Each utterance was normalised for loudness using the root mean square measure of the average power of the signal. This was done to remove any potential effect of stimulus loudness on learning. The model was trained on the first utterance by each speaker, and tested on the second. The training set was presented to the model ten times. The two digits were presented in an interleaved fashion. Informal tests demonstrated that on average the order in which the stimuli were presented did not significantly affect the performance of the trained models. It did, however, introduce higher trial to trial variability. Hence, we fixed the presentation schedule for the simulations described in this paper for a more fair model comparison. Each word was followed by 100 ms of silence. The silence was introduced to aleviate the confounding problem of having to perform word segmentation.

All reported results are calculated using model responses to the witheld test set of 188 distinct auditory stimuli (2 words spoken by the same 94 speakers as during training, but the particular pronunciations of the words were different from the training set). Each testing exemplar was presented 4 times, because due to the stochasticity of AN responses, input AN spike patterns in response to repeated presentations of the same word were not identical. This means that the results are reported in response to 752 testing examples in total (2 words * 94 speakers * 4 presentations).

### Spiking neural network architecture

To investigate whether information about auditory objects, such as words, is better encoded within spatio-temporal PGs rather than through rate encoding, we constructed a biologically grounded, unsupervised spiking neural network model of the auditory pathway, as shown in Fig. 1 (for full model parameters see Tbl. 1) [10]. The AN-CN-IC-CX model comprised of five layers: 1) the auditory nerve (AN); 2) the cochlear nucleus (CN), encompassing subpopulations representing three major ventral CN cell classes described by neurophysiologists: chopper (CH), onset (ON) and primary-like (PL); 3) the midbrain (inferior colliculus or IC) on which all CN subpopulations converge; and 4-5) cortical (CX) layers: primary (A1) and secondary (Belt) auditory cortex. We hypothesize that the subcortical layers (CN, IC) play a key role in reducing the physiological noise inherent to the AN. To investigate their importance we also constructed and evaluated a reduced AN-CX model without the CN or IC stages.

In the brain sub-populations of the CN do not necessarily synapse on the IC directly. Instead, they pass through a number of nuclei within the superior olivary complex (SOC). The nature of processing done within the SOC in terms of auditory object recognition (rather than sound localisation), however, is unclear. The information from the different CN sub-populations does converge in the IC eventually, and for the purposes of the current argument we model this convergence as direct. The same simplified connectivity pattern (direct CN-IC projections) was implemented by [13] for their model of the subcortical auditory brain.

Another simplification within our models has to do with the higher stages IC, A1 and Belt. They are not as neurophysiologically detailed as AN or CN. For example, the A1 stages of the full AN-CN-IC-CX and the reduced AN-CX models are supposed to be a loose and simplified approximation of the MGN and A1 in the real brain. Furthermore, we did not include recurrent connectivity in the cortical stages of our models. This simplification was done to be able to analyse the emergence and the nature of PGs for auditory stimulus encoding. Detecting PGs is a non-trivial problem even in feedforward architectures. It becomes even harder once recurrency is introduced. While we believe that recurrent connections are important for learning longer auditory sequences, and for dealing with speech in noise, we leave their inclusion for future work.

### Auditory Nerve (AN)

The AN comprised of 1000 medium spontaneous rate fibres modelled as described by [14], with log spaced characteristic frequencies (CF) between 300-3500 Hz, and a 60 dB threshold. The AN model by [14] has a high level of physiologically realism, ensuring that our models receive input which is highly representative of the signal processing challenges faced by the real auditory brain hierarchy. AN fibers, both biological ones and those of the model, are noisy channels plagued by “temporal and spatial jitter”. Temporal jitter arises when the propensity of the AN fibers to phase lock to temporal features of the stimulus is degraded by poisson-like noise in the nerve fibers and refractoriness [15]. “Spatial jitter” refers to the fact that neighbouring AN fibers have almost identical tuning properties, so that an action potential that might be expected at a particular fiber at a particular time may instead be observed in neighbouring fibers (the “volley principle” [16]). Both forms of jitter disrupt the firing pattern precision required for PG learning, and reducing the jitter should help the plastic CX layers of the model to learn the statistical structure of the stimulus set [10]. Jitter reduction can be accomplished by the CN and IC layers, which were modelled closely on anatomical and physiological characteristics of their biological counterparts [10].

### Neuron Model

Apart from the AN, which was modelled as described by [14], all other cells used in this paper were spiking neurons as specified by [17]. The spiking neuron model by [17] was chosen because it combines much of the biological realism of the Hodgkin-Huxley model with the computational efficiency of integrate-and-fire neurons. We implemented our models using the Brian simulator with a 0.1 ms simulation time step [18]. The native Brian exponential STDP learning rule with *nearest mixed (spike rule* and *STDP rule* parameters respectively in Tbl. 1) weight update paradigm was used [18]. A range of conduction delays (Δ_*ij*_) between layers is a key feature of our models. In real brains, these delays might be axonal, dendritic, synaptic or due to indirect connections [19], but in the model, for simplicity, all delays were implemented as axonal. The Δ_*ij*_ ∈ [0, 50] ms range was chosen to approximately match the range reported by [6].

**Excitatory Cells** Neurophysiological evidence suggests that many neurons in the subcortical auditory brain have high spiking thresholds and short temporal integration windows, thus acting more like coincidence detectors than rate integrators [3, 20]. This is similar to the behaviour of the Izhikevich’s Class 1 neurons [17]. All subcortical (CN, IC) excitatory cells were, therefore, implemented as Class 1. To take into account the tendency of neurons in the auditory cortex to show strong adaptation under continuous stimulation [21] Izhikevich’s Spike Frequency Adaptation neurons were chosen to model the excitatory cells in the auditory cortex (A1 and Belt).

**Inhibitory Cells** Since inhibitory interneurons are known to be common in most areas of the auditory brain [3, 21] except the AN, each stage of the models apart from the AN contained both excitatory and inhibitory neurons. Inhibitory cells were implemented as Izhikevich’s Phasic Bursting neurons [17]. Sparse connectivity between excitatory to inhibitory cells (*w^EI^_ij_*) within a model area was modelled using strong one-to-one connections from each excitatory cell to an inhibitory partner. Each inhibitory cell, in turn, was fully connected to all excitatory cells (*w^IE^_ij_*). Such an inhibitory architecture implemented dynamic and tightly balanced inhibition as described in [22], which resulted in competition between excitatory neurons, and also provided negative feedback to regulate the total level of firing within an area. Informal tests demonstrated that the exact implementation or choice of neuron type for within-layer inhibition did not have a significant impact on the results presented in this paper, as long as the implementation still achieved an appropriate level of within-layer competition and activity modulation.

### Cochlear Nucleus (CN)

The model CN was implemented as three parallel cell populations: 1000 CH neurons, 1000 PL neurons and 100 ON neurons. The CH cells removed space jitter while ON cells removed time jitter. Their activity was combined in the IC to produce spike rasters with reduced jitter in both space and time. The modelling choices for the three subpopulations of CN are described in more detail below.

In the brain CH cells receive a small number of afferent connections from AN neurons with similar CFs [23]. The incoming signals are integrated to produce regular spike trains. In the full AN-CN-IC-CX model, a CH subpopulation was simulated by units with Gaussian topological input connectivity, where each CH cell received afferents from a small tonotopic region of the AN (*σ* = 26 AN fibers). Their discharge properties correspond closely to those reported experimentally for biological CH neurons (Fig. 3, right column).

PL neurons make up ≈ 47% of the ventral CN in the brain [23], suggesting an important role in auditory processing. Although their contribution to the processing of AN discharge patterns is perhaps less clear, informal tests in our model indicate that their inclusion leads to significantly better model performance [10]. PL cells essentially transcribe AN firing [23] and were modelled using strong one-to-one afferent connections (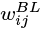) from the AN. The discharge properties of the model PL neurons also correspond closely to those reported experimentally (Fig. 3, left column).

ON cells are relatively rare, constituting around 10% of the ventral CN [23]. They require strong synchronized input from many AN fibers with a wide range of CFs in order to fire [20], which results in broad frequency tuning and enables them to phase-lock to the fundamental frequency (*F_0_*) of voiced speech sounds [26]. In the full AN-CN-IC-CX model, an ON cell population was simulated using sparse (34%) connectivity from across the AN. The interplay between converging ON and CH cell inputs to the IC can reduce jitter in the neural representation of vocalisation sounds.

**Fig 3.**
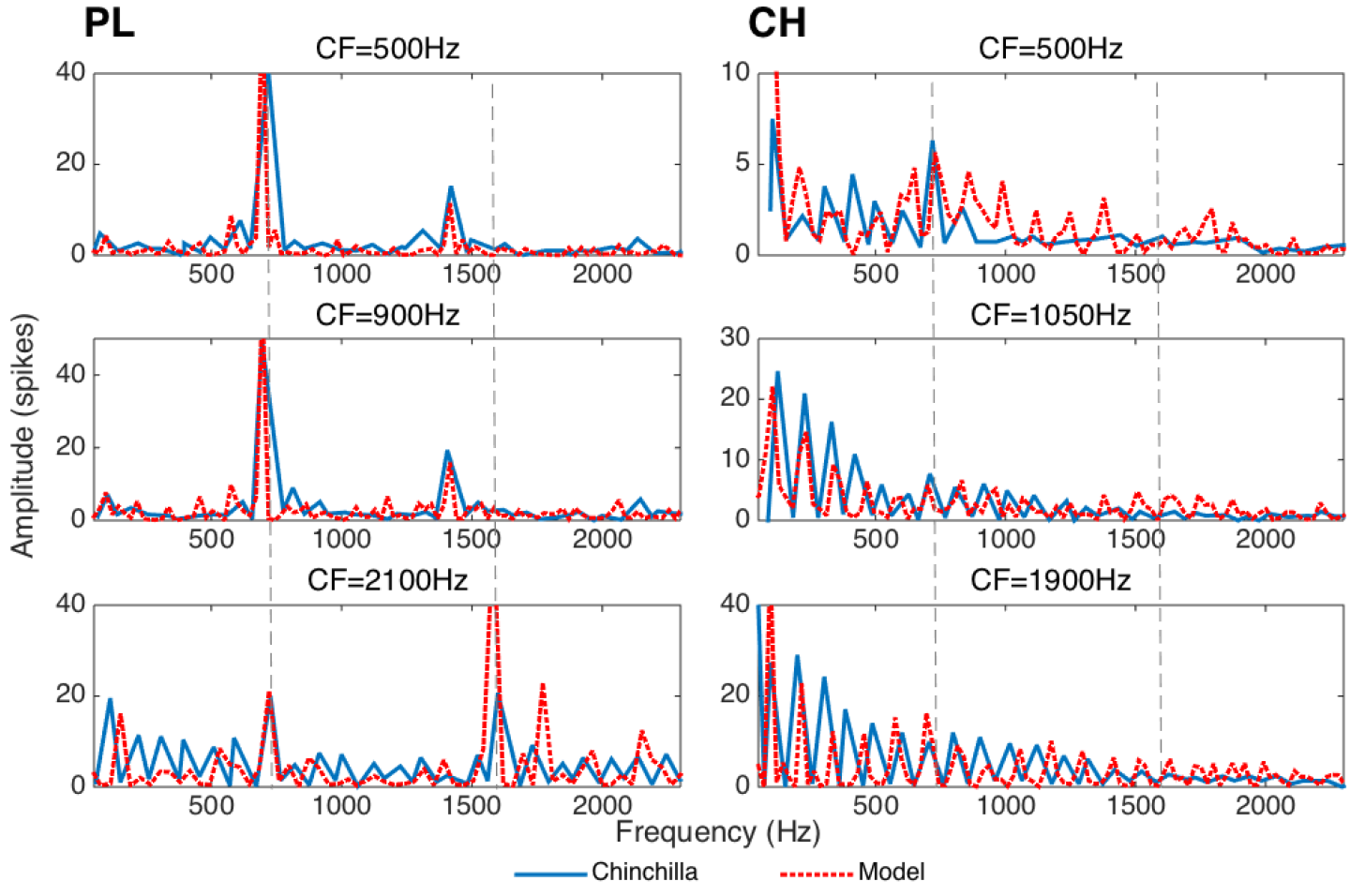
Spectra (computed as Fast Fourier Transforms of period histograms) of primary-like (PL) (left column) and chopper (CH) (right column) cochlear nucleus neuron responses to a synthetic vowel /a/ generated using the Klatt synthesiser [24]. The ordinate represents the level of phase-locking to the stimulus at frequencies shown along the abscissa. Dotted lines show the positions of the vowel formant frequencies *F_1_* and *F_2_*. Data from chinchilla CN fibers reproduced from [25] is shown in solid blue. Data collected from the corresponding model CN fibers is shown in dashed red. Similarity between the real and model fibers’ response properties suggests that the model’s performance is comparable to the neurophysiological data.

Since ON cells synchronise to the voice *F_0_*, they can introduce regularly spaced strong afferent input to the IC. Even if these afferent currents are sub-threshold, they nevertheless prime the postsynaptic IC cells to discharge at times corresponding to the cycles of stimulus *F*_0_. If IC cells also receive input from CH cells, then ON afferents will help synchronise CH inputs within the IC by increasing the likelihood of the IC cells firing at the beginning of each *F*_0_ cycle. This is similar to the neural coding ideas first described by [7].

### Inferior Colliculus (IC) and Auditory Cortex (CX)

Each model IC cell received one-to-one afferent connectivity from PL, narrow tonotopic connectivity from CH (*σ* = 2 neurons) and full connectivity from the ON cells. These subcortical connections (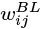) were not plastic. The IC→A1 and A1→Belt (and the equivalent AN→A1 and A1→Belt in the reduced AN-CX model) connectivity was full, feedforward, with STDP learning [8] (parameters *α_p_*, *α_d_*, *τ_p_* and *τ_d_* in our models, see Tbl. 1) and a uniform distribution of conduction delays (Δ_*ij*_ ∈ [0, 50] ms) [27]. The initial afferent connection strengths were randomly initialised using values drawn from a uniform distribution 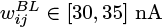.

### Polychronization index

As outlined in the introduction, we envisaged that our model of the auditory brain would exhibit unsupervised learning of speaker independent representations of naturally spoken words “one” and “two” by forming a hierarchy of stable spatio-temporal patterns of firing (PGs). An exhaustive search for PGs through the network was prohibitive, especially given that the number of neurons participating in each PG was unknown a priori, and could be large and variable [28]. We therefore developed a numerical score, the “polychronization index” (PI) to quantify the prevalence of reproducible patterns of firing across the population of neurons in one layer.

The PI was calculated for each cell *j* within a particular stage of the model. For each spike produced by cell *j* we searched for spikes fired by other cells within the same stage of the model within a fixed time interval Δ*t_j_*. PGs which are informative of a stimulus class should be reproducible across the different presentations of different exemplars *e_k_*(*s*) of the stimulus class, where *k* ∈ {1,…, 376} is the number of exemplars (4 repetitions of 94 pronunciations) of each of the s = 2 stimulus classes (words “one” and “two”). For each presentation of stimulus exemplar *e_k_*(*s*) we randomly selected one spike by cell *j* and noted its time *t_j_* post stimulus onset. We then constructed a 1000 neuron by 101 ms matrix 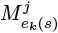 representing the firing of the other neurons within the same stage of the model that occured within Δ_*t*_ = *t_j_* ± 50 ms at 1 ms resolution. The ±50 ms time window reflects the maximum conduction delay (Δ_*ij*_) in our model, which puts an upper limit on the temporal integration window of neuron *j*. If neuron *j* is part of a PG selective of a particular stimulus class *s*, then we would expect to see similar firing pattern matrices 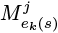 for different *e_k_*(*s*). Consequently, elements of 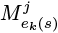 which are non-zero across the different stimulus exemplars more frequently than would be expected by chance (where chance levels can be estimated from the average firing rate *f*) are diagnostic of the PG firing pattern. The larger the proportion of stimulus exemplars for which these elements are non-zero, the more established and reproducible the PG firing pattern is across the responses to the different exemplars from the given stimulus category.

We therefore computed 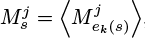, where 〈·〉 signifies the mean over all the exemplars *e_k_(s)* of stimulus class s, and then identified the ten elements of 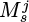 with the largest mean spike counts 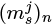, where *n* ∈ {1,…, 10} (the element corresponding to the randomly sampled spike by cell *j* occurring at time *t* was ignored). These were used to compute 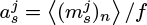, where *f* is the average firing rate within the layer and 〈·〉 signifies the mean over the ten largest elements indexed by *n*.

Thus, *a_s_^j^* quantifies the evidence that cell *j* participates in polychronous firing in its responses to stimulus class s. To calculate an overall polychronization index which is not stimulus specific we simply compute 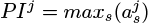. The larger the *PI*^*j*^, the stronger the evidence that cell *j* takes part in a PG.

### Information analysis

Apart from quantifying whether PGs arise throughout the hierarchy of our model of the auditory brain, it is imporant to measure how informative these PGs are about the two stimulus classes, words “one” and “two”. One common way to quantify such learning success is to estimate the mutual information between stimulus class and neural response *I*(*S*;*R*). It is calculated as 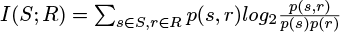, where *S* is the set of all stimuli and *R* is the set of all possible PG responses, *p(S, R)* is the joint probability distribution of stimuli and responses, and 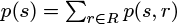 and 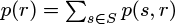 are the marginal distributions [29]. Stimulus-response confusion matrices were constructed using decoder multi-layer perceptron (MLP) networks (see below), and used to calculate *I*(*S*; *R*).

Our information analysis approach uses observed frequencies as estimators for underlying probabilities *p*(*s*),*p*(*r*) and *p*(*s,r*). This introduces a positive bias to our information estimates 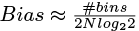, where *#bins* is the number of potentially non-zero cells in the joint probability table, and *N* is the number of recording trials [29]. Given the large *N* in our tests of model performance (*N* = 752), the bias was negligible (*Bias* ≈ 0.004 bits) and was ignored.

The upper limit of mutual information to be achieved by PGs in our models *I*(*S*; *R*) is given 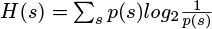, which, given that we had two equiprobable stimulus classes, here equals 1 bit.

**Using multi-layer perceptron (MLP) decoders to evaluate network performance:** Decoder MLPs were used to evaluate the ability of PGs within the full AN-CN-IC-CX and the reduced AN-CX models of the auditory brain to represent stimulus identity. We compared the amount of information about the two stimulus classes “one” and “two” when using either rate or temporal encoding schemes in the two models. The MLPs were trained to classify 752 input vectors *x*_*i*_ ∈ *R*^201^ as either word “one” or “two”. The nature of the input feature vectors *x_i_* for the temporal and rate encoding schemes is described below.

Our MLP decoders had a small, single hidden layer of 20 units (10% of the input layer size) to limit their capacity and therefore to make them more sensitive to the informativeness of different auditory brain models and encoding schemes under investigation. The MLPs had hidden layer neurons with hyperbolic tangent transfer functions. They were trained until convergence using scaled conjugate gradient descent [30] and a cross-entropy loss function. Once the MLPs were trained, their classification outputs were used to fill the confusion matrices used in the information analysis calculations described above. The process was repeated 20 times, each time with a different random subsample of *J* cells, to obtain a distribution of stimulus information estimates for each cell population.

**Temporal code**: Temporal encoding assumes that information about stimulus class is provided by spatio-temporal firing patterns (PGs). In order to be informative, the same PG must be present more frequently in response to the exemplars of one stimulus class than the other. The vectors *x_i_* used as input to the decoder MLPs were designed to capture PGs within a particular stage of the auditory brain model. As discussed previously, it is unfeasible to explicitly identify PGs in our models. We do not know which neurons within a stage of the auditory brain model participate in a PG and which ones do not. We therefore introduce an approximate computationally feasible protocol to get an indication of the presence of PGs within a stage of the model, and then to use these approximations to calculate how informative the approximated PGs in this stage are. We appreciate that our meaure is not perfect, but we accept it, because it approximately lowerbounds the informativeness of PGs in the models, and it is informative enough to differentiate between the different models.

Our protocol takes the following form. First, in order to restrict the computational load, we randomly sample *J* = 100 cells from within the chosen auditory model stage. We make an assumption that every cell *j* within this sample takes part in a PG. We then train a separate MLP_*j*_ to try and classify the two stimulus classes, words “one” and “two”, using the approximated PGs that each cell *j* is part of. If our assumption was correct and cell *j* indeed participated in a PG, then MLP_*j*_ will achieve high classification accuracy, otherwise the classification accuracy will be low.

We quantified reproducible PG spatio-temporal patterns for each cell *j* using methods analogous to those used to calculate the polychronization index (PI) (see Sec. Polychronization Index). For each cell *j* we first computed a 100 neuron by 101 ms matrix 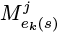. This was computed in the same was as for the PI score (see Sec. Polychronization Index), where 100 neurons are the randomly sampled subset of 100 neurons within the auditory model stage. For each of the 752 input stimuli *e*_*k*_(*s*) we then concatenated together a 100 element vector of row sums and a 101 element vector of column sums of matrix 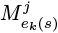 to form the 201 element column vectors *x_i_* for training the MLP_*j*_. Summing and concatenating reduces the dimensionality of the feature vectors from 100 * 101 = 10,100 (the dimensionality of matrix 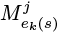)) to 201, while still capturing the key patterns of firing across the sample population of 100 cells in response to a single presentation of stimulus *e*_*k*_(*s*) and the amount of spatio-temporal structure in the network activity that would be compatible with the presence of PGs. Each of the decoders MLP_*j*_ was then trained to classify response vectors *x_i_* as either the word “one” or word “two”.

The confusion matrix used in the information analysis calculations described above was constructed based on the majority classification votes among the 100 trained MLP_*j*_ in response to each stimulus presentation.

**Rate code**: A random subset of *J* = 201 cells within an auditory brain model stage was chosen to match the dimensionality of the input vectors *x_i_* for training the MLPs for the temporal code informativeness estimation. The average firing rate of the 201 subsampled neurons was recorded in response to each of the 752 stimulus presentations to form the input to the rate code MLP.

## Results

In this paper we propose that speaker-independent word representations are encoded by unique discriminative spatio-temporal patterns of firing (PGs) in the auditory cortex. We test this hypothesis using a biologically inspired spiking neural network model of the auditory brain. Since we cannot explicitly detect the information bearing PGs due to computational restrictions, we present instead multiple pieces of evidence for the emergence of informative PGs within the plastic cortical stages of our model. These pieces of evidence include the particular change in the distribution of connection weights (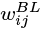) after training that is characteristic of PG-based learning, the increased polychrony of firing in the final cortical stages of the model (measured by the polychronisation index we devised in Sec. Polychronization Index), and the performance of MLP decoders trained to be sensitive to the existence of stimulus class selective PGs. The observed differences according to these measures between the full AN-CN-IC-CX and the reduced AN-CX models, and between rate and temporal encoding schemes, provide evidence in support of our hypothesis that more information is carried using temporal PGs rather than rate codes, and that the emergence of such informative PGs is only possible if stable input firing patterns are provided to the plastic cortical stages of our models.

**Changes in synaptic weights resulting from unsupervised learning**: To examine the synaptic weight changes that occurred in the cortical stages of the full AN-CN-IC-CX model during training we plotted the distributions of IC→A1 and A1→Belt connections (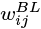) before and after training (Fig. 4, top row). While the majority of weights grew weaker, some connections were strengthened. Such a pattern of change is characteristic of learning PGs. Since the STDP configuration in our model is reminiscent of that described in [5], the majority of connections (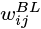) weakened due the non-informative random firing in the input. Some connections strengthen due to the presence of repeating stable patterns of firing (PGs) among the pre-synaptic neurons.

Compare this pattern of connectivity change to the equivalent stages of the reduced AN-CX model. Almost all AN→A1 and A1→Belt synaptic weights were weakened during training (Fig. 4, bottom row). In the reduced model, stochastic jitter in the AN presumably scrambled the structure of input patterns sufficiently to prevent regularities (PGs) in the inputs to be discovered and learned.

**Fig 4.**
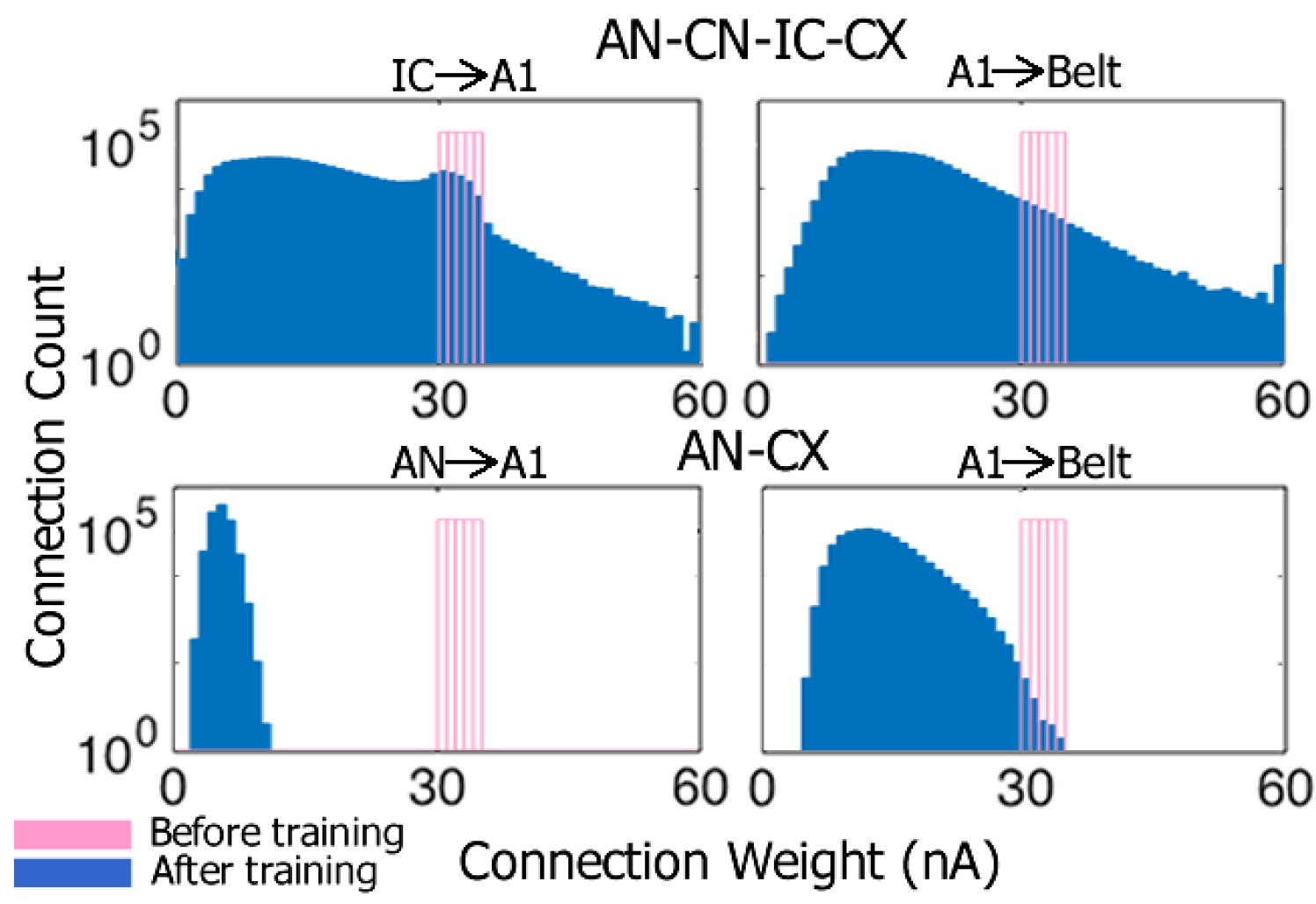
Distributions of synaptic connection weights (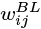) before (pink) and after (blue) training for the full AN-CN-IC-CX model (top row) and reduced AN-CX model (bottom row). The ordinate is log scaled.

These data suggest that PG learning relies on stable patterns of firing that serve as input to the plastic cortical stages A1 and Belt of the models. AN stochasticity hinders such learning, but de-noising of AN firing within the CN and IC makes PG-based learning possible again in the cortical stages of the full AN-CN-IC-CX model.

**Denoising of AN firing patterns and the emergence of polychronous groups (PGs)**: Fig. 5 shows that subcortical preprocessing in the CN and IC led to more stable spatio-temporal discharge patterns, as evidenced by higher PI scores in the IC compared to AN. These more reproducible firing patterns also carried through to A1 and Belt. The AN-CX model, which lacked subcortical preprocessing layers CN and IC, did not achieve the same stability of firing patterns seen in IC and CX of the full model even after training.

**Polychronous groups**: Fig. 6 shows a partial visualisation of three PGs that emerged in the A1 layer of the trained AN-CN-IC-CX model. These PGs responded preferentially either to different pronunciations of the word “one” or “two”. The red dots show the ten elements of the mean firing pattern matrix 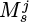 with the largest mean spike counts 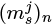 (see Sec. Polychronization Index). They can be thought of as partial PGs observed in response to the respective stimulus classes. When projected through the A1→Belt connections (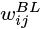) and the corresponding axonal delays (Δ_*ij*_), these patterns produce near-synchronous input from four or more A1 neurons in a small subset of Belt neurons, and these inputs are consequently strengthened during training. The green and yellow dots show such inputs for two Belt neurons which in this manner became selective for the word “one”, the white dots show equivalent data for a Belt neuron that became selective for “two”.

**Fig 5.**
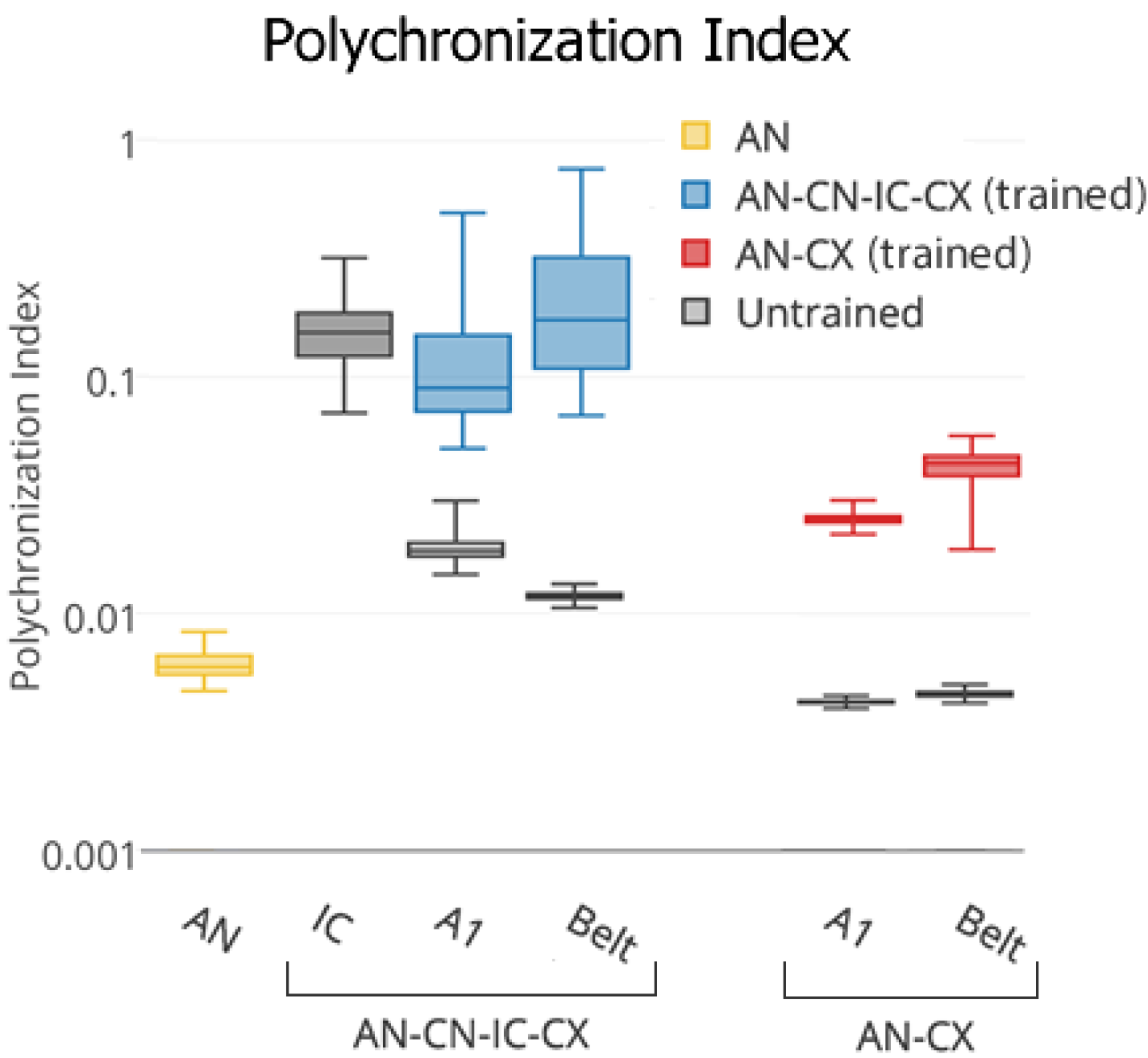
Box-and-whisker plot showing the distribution of polychronization indices over all the cells within the relevant layers of the full AN-CN-IC-CX and reduced AN-CX models. The ordinate is log scaled.

**Stimulus category encoding across models and layers**: Fig. 7 shows estimates of the speech stimulus identity information encoded in various layers of the full AN-CN-IC-CX and the reduced AN-CX models. Note the different y-scale ranges in panels A and B. Temporal encoding provides substantially more stimulus category information than rate encoding at every stage of the model. After training, the responses of the Belt layer of the full model carried as much as 0.52 bits per response, i.e. they could be decoded with an accuracy of 89.36% correct (672/752 correct trials). In comparison, in the untrained model, the stimulus category information never exceeded 0.05 bits (62.77% correct).

**Fig 6.**
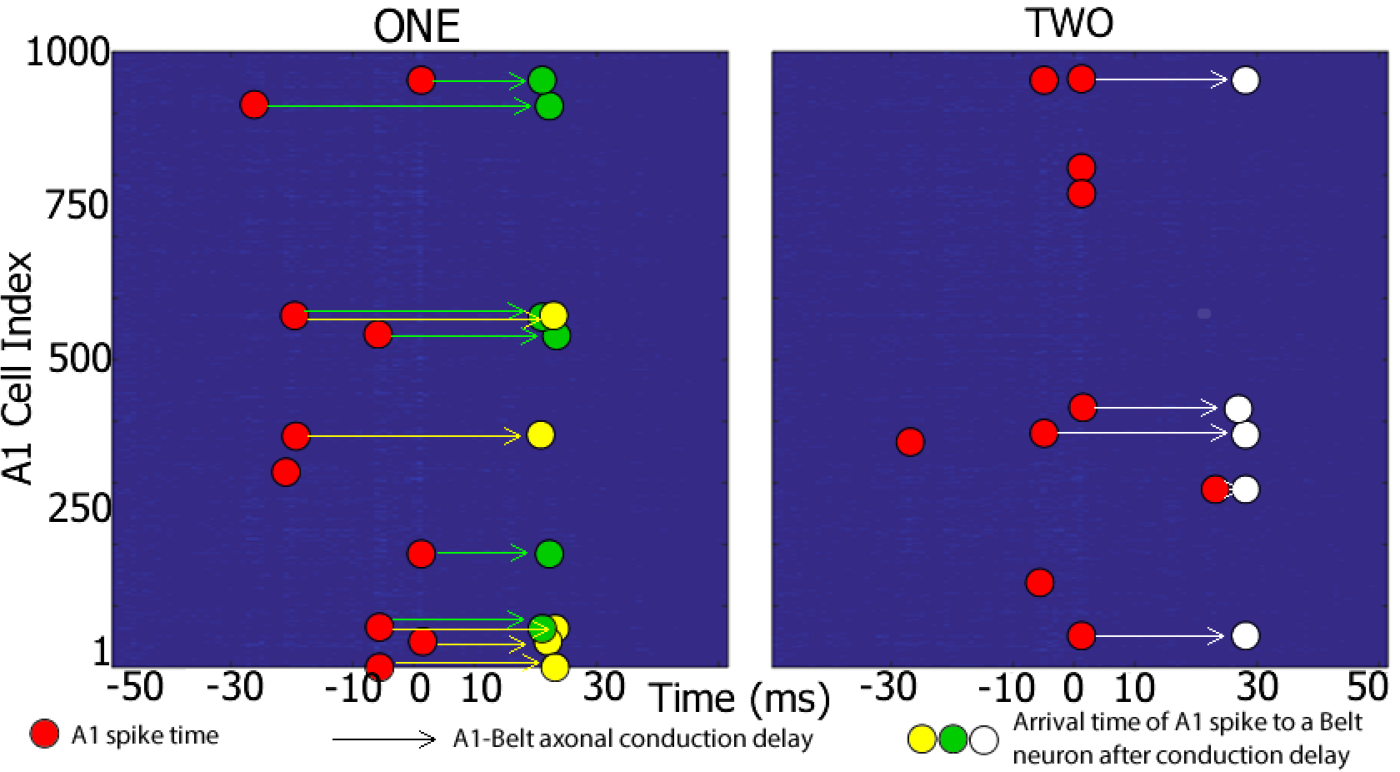
Evidence for PGs responding selectively to word “one” or “two” in the A1 layer of the trained AN-CN-IC-CX model. Each plot shows an example of a stable spatio-temporal spike pattern in A1 (red circles) in response to different pronunciations of the words “one” (left) and “two” (right). These spikes take part in at least one polychronous group that is selective for the particular word. In other words, these patterns are more likely to appear when an example of their preferred word is pronounced compared to an example of a non-preferred word. When projected through the A1→Belt connections 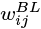) with different conduction delays (Δ_*ij*_) (arrows), these patterns produce near-synchronous input from several A1 neurons onto a subset of Belt neurons (green, yellow or white circles corresponding to three separate Belt neurons with different distributions of axonal conduction delays Δ_*ij*_). The green and yellow circles show such inputs for two Belt neurons which in this manner respond selectively for a number of different pronunciations of the word “one”, the white circles show inputs for a neuron that responded selectively to exemplars of the word “two”. Abscissa represents the time window Δ_*t*_ = *t_j_* ± 50 ms around the origin. The origin is centered around all the times *t* when a chosen A1 neuron *j* fires (see Sec. Polychronization Index for details). Ordinate represents the 1000 neurons that make up A1 in the AN-CN-IC-CX model. Red circles show the ten elements of the firing pattern matrix 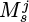 with the largest mean spike counts 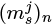 (see Sec. Polychronization Index for details).

A control simulation was run to test the ability of an untrained AN-CN-IC-CX model with the same distribution of IC-A1 and A1-Belt afferent connection weights (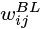) as the trained model to discriminate between the two stimulus classes. This was done by randomly shuffling the corresponding connection weights (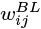) of the trained AN-CN-IC-CX model before presenting it with the two naturally spoken word stimuli.

**Fig 7.**
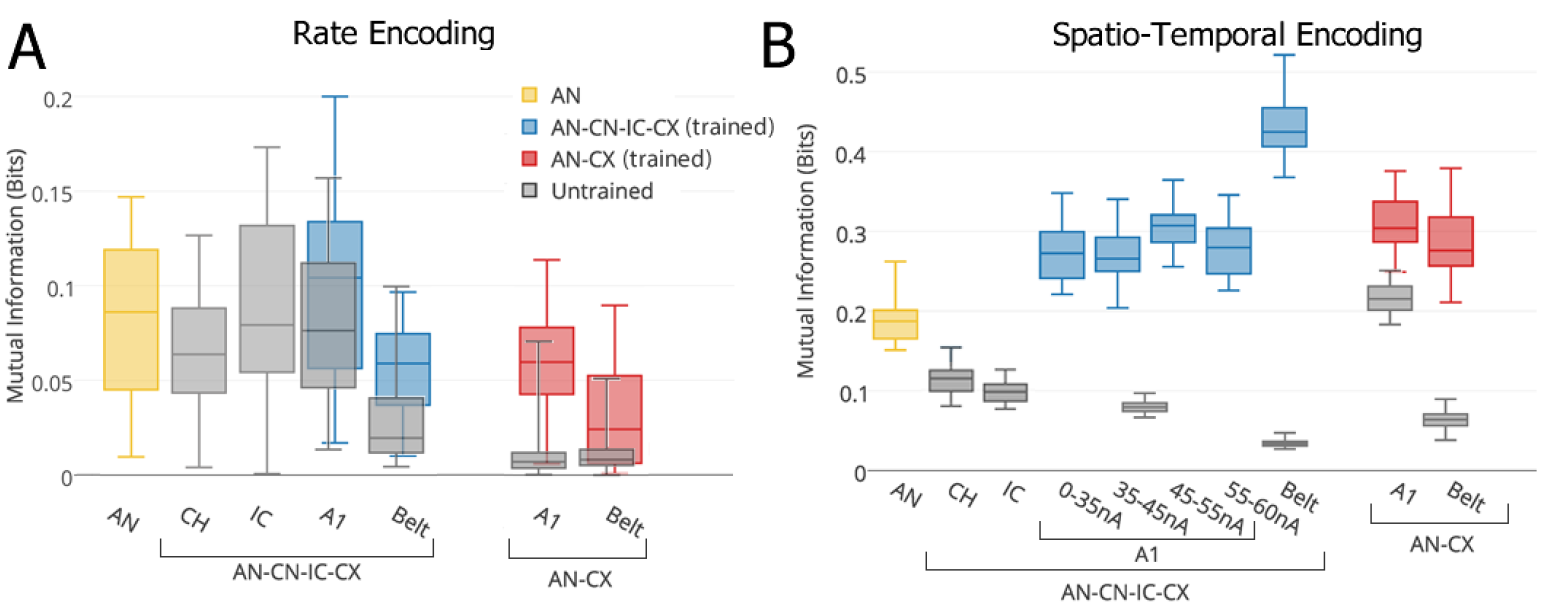
Box-and-whisker plots showing the distribution of information over twenty different subsamples of cells within various layers of the full AN-CN-IC-CX and reduced AN-CX models based on rate (**A**) and temporal (**B**) encoding schemes. In **B**, the A1 neurons are subdivided according to the strength of their connections (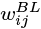) to the Belt layer. PL and ON layers are not shown. The ON layer conveys 0 bits of information, and PL is equivalent to AN.

It was found that such a model performed at the same level as the untrained model with afferent connection weights initialised from the 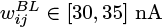 uniform distribution (0.05 (shuffled) vs 0.08 (random) bits in the A1, and 0.04 (shuffled) vs 0.03 (random) bits in Belt).

Fig. 7B also shows that responses in the plastic A1 and Belt layers of the full AN-CN-IC-CX model contain significantly more stimulus category information after unsupervised learning than the input AN layer responses. This indicates that the biologically inspired spiking AN-CN-IC-CX model has learnt to develop a more efficient, less redundant and more informative representation of the naturally spoken word stimuli during training, in line with [31]. This is achieved using only physiologically realistic, local STDP learning. After learning, the Belt area of the full AN-CN-IC-CX model encoded more than twice as much stimulus category information as the AN, but it failed to reach the maximum of 1 bit of information required for perfect word identification. The information encoding performance of the AN-CN-IC-CX model should be possible to improve by the addition of recurrent plastic within-layer cortical connections or additional plastic cortical layers. We leave this, however, for future work.

When the trained model was tested on additional data of the same words “one” and “two” being spoken by twenty novel speakers (ten male and ten female speakers pronouncing each word twice), the network reached a similar level of performance as described above (0.27 (new data) vs 0.28 (old data) bits in the A1, and 0.48 (new data) vs 0.43 (old data) bits in Belt).

It is interesting to note that those A1 cells which acquired A1→Belt connections in the 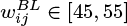 range are more informative than A1 cells with maximally strengthened connections. This effect is in line with the work by [5]. In order to maximally strengthen the A1→Belt connections (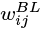), PGs in the A1 have to appear very freuquently. This is more likely to happen if the PGs are present in response to both stimulus classes “one” and “two”. Such PGs, however, are not informative of the stimulus class identity. This effect also explains why A1 and Belt stages of the full AN-CN-IC-CX model have higher levels of spatio-temporal stimulus category information than the IC stage of the model despite no increase in the PI from IC upwards. This is because PI score only measures the presence of PGs in a particular stage of the model without measuring their informativeness. The data suggests that while the degree of stability of spatio-temporal firing in the IC, A1 and Belt stages of the full AN-CN-IC-CX model is similar, the stimulus class selectivity and hence the informativness of PGs grows throughout this feedforward hierarchy.

## Discussion

In this paper we argued that a hierarchy of speaker-independent informative PGs is learnt within the different stages of the plastic cortical layers of the full AN-CN-IC-CX model. The learning in the model, however, is reliant on the input of stable firing patterns to the plastic cortical stages A1 and Belt. Such stable firing patterns are obscured by stochasticity in the raw AN firing rasters [10]. Consequently the cortical layers are essentially unable to learn speaker independent representations of naturally spoken words using unprocessed AN input (reduced AN-CX model). Subcortical preprocessing in the CN and IC stabilises and de-noises the AN firing patterns, thus allowing the cortical ensembles of the full AN-CN-IC-CX model to form category specific response patterns. The biological realism of the inputs to our model sets our results apart from the similar work by [32], who showed that a recurrent network of winner-take-all microcircuits with STDP learning is capable of achieving similar informativeness for differentiating between words “one” and “two” (albeit for only three utterances pronounced by two speakers) as was achieved in our model (around 0.6 bits). They also argued for temporal information encoding.

We took inspiration from the known neurophysiology of the auditory brain in order to construct the spiking neural network models used in this paper. As with any model, a number of simplifying assumptions had to be made with regards to certain aspects that we believed were not crucial for testing our hypothesis. These simplifications included the lack of superior olivary complex or thalamus in our full AN-CN-IC-CX model, the nature of implementation of within-layer inhibition in both the AN-CX and AN-CN-IC-CX models, and lack of top-down or recurrent connectivity in either model. While we believe that all of these simplifications do affect the learning of auditory object categories to some extent, we also believe that their particular implementation in our models does not undermine out qualitative findings and conclusions.

The full AN-CN-IC-CX model of the auditory brain described in this paper possesses a unique combination of components necessary to simulate the emergent neurodynamics of auditory categorisation learning in the brain, such as biologically accurate spiking dynamics of individual neurons, axonal conduction delays, STDP learning, neuroanatomically inspired architecture and exposure to realistic speech input. With its biological realism, the full AN-CN-IC-CX model described in this paper can be used to make testable predictions about the auditory object encoding in the auditory brain. For example, the reduction of jitter in spiking responses as one ascends from AN through IC to auditory cortex should be observable in physiological experiments. Furthermore, large channel count recordings may make it possible to discover PG firing patterns which encode categorical stimulus information in the activity of ensembles of real cortical neurons.

## Acknowledgments

This research was funded by a BBSRC CASE studentship award (No. BB/H016287/1), and by the Oxford Foundation for Theoretical Neuroscience and Artificial Intelligence.

